# Sigma factors as potential targets to enhance recombinant protein expression

**DOI:** 10.1101/2024.11.12.623148

**Authors:** Laura Pohlen, Emily García, Luz María Martínez, Noemí Flores, Jochen Büchs, Guillermo Gosset, Alvaro R. Lara

## Abstract

The transcriptional factors control the expression of many genes and represent an important layer of complexity in cell factories. However, the effect of individual sigma factor deletions from a biomanufacturing perspective has not been addressed. In this contribution, growth and oxygen consumption of *E. coli* BW25113 strains with individual inactivation of each sigma factor were characterized under various conditions. The deletion of *rpoD* has been reported to be lethal for *E. coli*. However, the evaluated *rpoD* mutant did not exhibit major growth defects in the mineral medium and is explained because the *rpoD* gene is duplicated in this strain. Recombinant protein was expressed at three different induction levels in a mineral and LB media. The *fliA* mutant was the best producer in the mineral medium, whereas the *rpoD* mutant overperformed the other strains in LB medium. This suggest that a lower *rpoD* gene dosage is positive for the performance of the cell factory in complex medium. In cultures at 20 °C, the *rpoS* mutant exhibited the greatest recombinant expression. To our knowledge, this is the first systematic study evaluating the potential of sigma factor deletion for improving recombinant protein production.

---

Bacterial RNA polymerases require subunits, named sigma (σ) factors to recognize specific promoters and start the transcription process. In *Escherichia coli*, the “housekeeping” and essential σ^70^(*rpoD*) controls the expression of many genes expressed during exponential growth (Shimada et al., 2014). Alternative σ-factors have been described in *E. coli*, σ^54^ (*rpoN*), σ^38^ (*rpoS*), σ^32^ (*rpoH*), σ^28^(*fliA*), σ^24^(*rpoE*, which is an essential gene), and σ^19^ (*fecI*). Cho and coworkers reconstructed a genome-scale network of the σ-factors and reported 4,724 σ-factor specific promoters (Cho et al., 2014). While the σ-factors control the expression of many genes, whether such genes are dispensable for the cell factory under production conditions remains largely unstudied, but there are some reports suggesting that σ-factor deletions can be useful to improve cellular performance. For instance, it has been shown that the deletion of *rpoS* increases the production of recombinant protein in chemostat cultures of *E. coli* at low dilution rates (Chou et al., 1996). Bafna-Rührer and coworkers cultured *E. coli rpoS* and *fliA* mutants under short periods of glucose starvation and found that the *rpoS* deletion improved the biomass yield, but *fliA* deletion worsened it (Bafna-Rührer et al., 2024). Despite the physiological relevance of the σ-factors, there is not, to the best of our knowledge, a systematic study on the effect of individual knockouts on the growth and expression of recombinant protein under different culture conditions. Here, the performance of the individual σ-factors mutants was characterized in a mineral medium and in lysogeny broth (LB), at 20 and 37 °C in microbioreactors with online measurement of the oxygen transfer rate (OTR), biomass and NADH or green fluorescence protein fluorescence signals.

The impact of individual sigma factor deletion was first evaluated in a buffered mineral medium supplemented with 4 g L^-1^ glucose. The strains were cultured at 37 °C, except for the *rpoH* mutant, which did not grow at this temperature. The growth profiles for the strains were similar, exhibiting a clear relation between biomass accumulation, OTR and NADH fluorescence (Fig. 1a). Notwithstanding, the NADH fluorescence remained constant during the stationary phase for all strains except for the *rpoS* mutant, in which a slow but steady increase was observed until 18 h of culture. Possibly the lack of control of genes related to the stationary phase promoted some metabolic activity using intracellularly available metabolite pools, affecting the redox state of the cell. Apart from the *fliA* and *fecI* mutants, which grew at the same rate as the WT, the other sigma factors deletions decreased the growth rate between 13 and 30 % (Fig. 1b). The scattered light signal indicated that the biomass synthesis was similar for all the strains, with small differences for *rpoN* and *rpoS* mutants (Fig. 1b). The specific NADH fluorescence intensity was 25 % lower for the *rpoD* and *rpoS* mutants, which suggests a decreased energy metabolism in these genotypes. The accumulated oxygen transferred to the liquid was lower for all the mutants except *rpoN (*Fig. 1b). This can be related to decreased protein formation in the mutants, which results in ATP savings. In turn, deletion of the *rpoN* gene affects nitrogen metabolism by limiting the uptake of ammonium into the cell, which may explain the reduced growth of this mutant. Thus, the *rpoN* mutants may perceive nitrogen limitation, increasing the activity of the glutamine synthetase, which increases the ATP demand by 14 %, compared to nitrogen-excess conditions (Reitzer and Schneider, 2001). This may contribute to an oxygen demand similar to that of the wild-type, and higher than the rest of the mutants. Interestingly, the accumulated oxygen transfer rate for the *fecI* and *fliA* mutants decreased by 18 and 11 %, respectively, compared to the WT strain, despite all three exhibiting similar growth rates (Fig. 1b). The lower oxygen consumption by the *fliA* mutant is probably consequence of the absence of flagellum synthesis and flagellum rotation, which are highly energy-demanding processes (Schavemaker and Lynch 2022). This may result in less energy being required and thus less oxygen consumed by the cells. However, the reasons for reduced oxygen consumption by the *fecI* mutant remain elusive.

**Figure 1.**
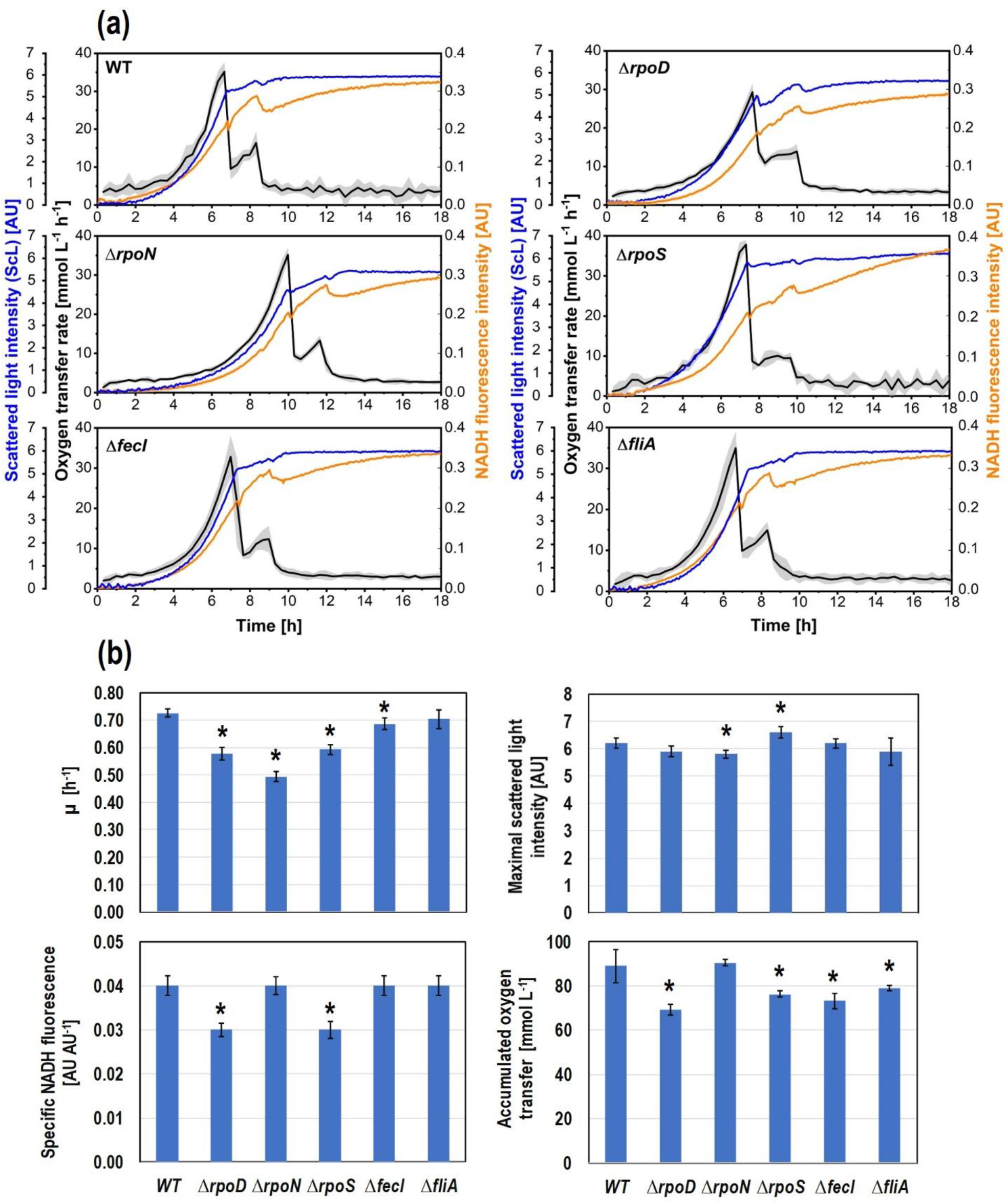
(a) Culture profile of the untransformed strains in mineral medium. Cell growth was monitored as scattered light (ScL) intensity. The oxygen transfer rate and NADH fluorescence were also monitored. Shaded bands indicate the standard deviation between replicates. (b) Main culture parameters. Specific growth rate (*µ*) and NADH fluorescence were calculated over the exponential growth phase. * Indicates significant difference (*n* = 4; *p*<0.05), compared to the WT strain.

The growth of the *rpoD* mutant was unexpected, since σ ^70^is considered an essential gene (Goodall et al., 2018). Examination of the mutant genome revealed the presence of a second copy of the *rpoD* gene (Fig. S1), in agreement with a previous report for strain BW25113 (Yamamoto et al., 2009). The *rpoD* expression, relative to the housekeeping gene *ihfB* was determined for the mutant and WT strains by RT-qPCR. The results are depicted in Fig. S2. The *rpoD* expression level in the mutant strain was approx. 20 % lower than in the WT (*n* =7, *p*<0.07). The expression of the duplicate *rpoD* appears to be insufficient to retain the growth rate at the same value as the WT in the mineral medium (Fig 1b).

The expression of recombinant protein was tested in the mineral medium within all strains using GFP as a model. The *gfp* gene was coded in a high copy-number plasmid under transcriptional control of the *lac* promoter. GFP expression was induced with 0.1, 0.2 or 0.3 mM IPTG. Culture profiles are shown in Fig. 2. For a given strain, a higher GFP fluorescence is related to a lower maximal OTR (Fig. 2a), which can be connected to a higher metabolic burden (Ladner et al., 2017; Mülmann et al., 2018). The growth rate of the WT and *rpoS* mutant strains was not affected by the induction level (Fig 2B). In contrast, the growth rate of the other mutant strains using 0.3 mM IPTG was lower than the corresponding value at 0.1 mM IPTG (Fig 2B). In comparison to the WT strain, the growth rates of the *rpoD, rpoS* and *fliA* mutants were considerably lower. The accumulated biomass was affected slightly, showing only small decreases for mutants other than *rpoN* (Fig. 2B). Remarkably, the deletion of sigma factor genes resulted in strongly decreased GFP expression, compared to the WT, being *fliA* the only exception. The *fliA* mutant attained approx. 3 and 4-fold higher maximal and specific GFP fluorescence, respectively, than the WT strain. Moreover, despite the growth rate reduction of the *fliA* mutant, the specific GFP fluorescence emission rate in cultures of this mutant was between 1.8-3-fold greater than for the WT strain (Fig. 2b). When induced with 0.2 and 0.3 mM IPTG, all the mutants exhibited higher accumulated oxygen transfer rates compared to the corresponding values of the WT (Fig. 2b). Interestingly, the greatest accumulated oxygen transfer rate was seen in cultures of the *rpoS* mutant induced with 0.2 mM IPTG, even though under such conditions, this strain grew slower, produced less biomass, and expressed less GFP than the other strains (Fig. 2b). This results in higher stoichiometric oxygen demand to oxidize glucose to CO_2_ for cellular maintenance during the longer fermentation times of this strain. In contrast, in cultures of the *fliA* mutant induced with 0.2 mM IPTG, the accumulated oxygen transfer was the lowest among all the mutants, while the GFP specific fluorescence and emission rate was the highest (Fig. 2b).

**Figure 2.**
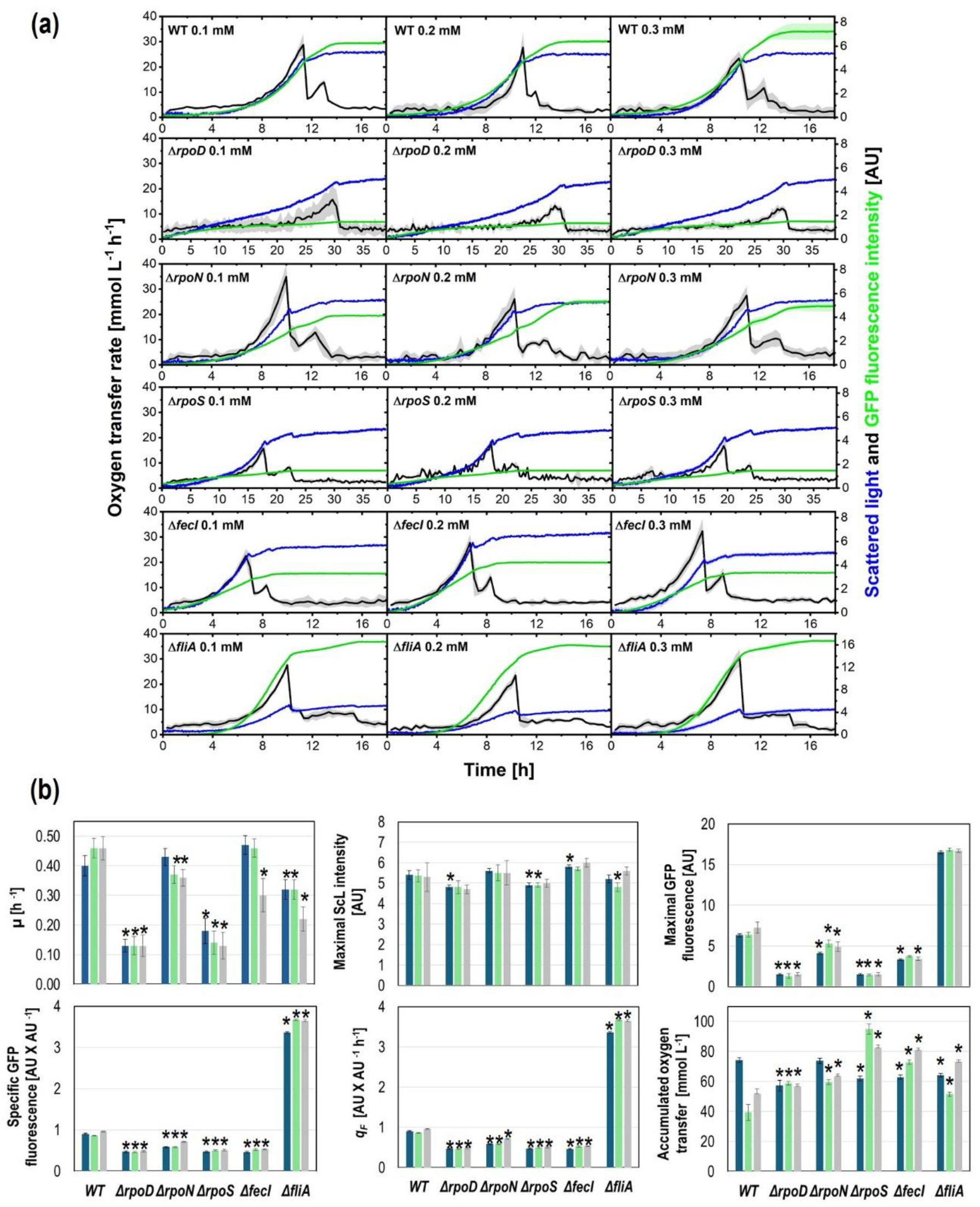
(a) Culture profile of the strains bearing the plasmid pV21 in mineral medium at 37 °C. GFP expression was induced with 0.1, 0.2 or 0.3 mM IPTG from inoculation. Cell growth was monitored as scattered light (ScL) intensity. The oxygen transfer rate and GFP fluorescence were also monitored. Shaded bands indicate the standard deviation between replicates. (b) Main culture parameters using 0.1 (blue bars), 0.2 (green bars), or 0.3 (grey bars) mM IPTG. Specific growth rate (*µ*), GFP fluorescence, and GFP fluorescence emission rate (*q*_*F*_) were calculated over the exponential growth phase. * Indicates significant difference (*n* = 4; *p*<0.05), compared to the WT strain under the same conditions.

There is no clear relationship between the observed parameters. In general, the effects were more evident for the *rpoS* and *fliA* mutant strains. The positive effect observed in the *fliA* mutant can be attributed to better resource stewardship, due to less energy spent in motility-related protein synthesis. Similar effects have been observed by deleting genes coding for the transcriptional dual regulator FlhC, which is the main regulator of flagellum synthesis and swarming migration (Han et al., 2021; Lara et al., 2024). Ziegler and coworkers also reported an increase in the production of a recombinant protein lacking *fliA* and some operons related to motility (Ziegler et al., 2021). In the case of the *rpoS* mutant, it is known that flagellar synthesis is upregulated, when the *rpoS* gene is deleted (Dong and Schellhorn 2009), which may partially explain the growth rate reduction and accumulated oxygen transfer increase.

Recombinant protein expression at low temperatures can increase the production of correctly folded proteins in *E. coli* (San-Miguel et al., 2013). The use of expression systems engineered to induce protein expression in response to low temperature, is attractive to attain high expression levels (Lin et al., 2024). Therefore, a set of cultures at 20 °C in the mineral medium was carried out, which included the *rpoH* mutant. As explained above, increasing the IPTG concentration from 0.2 mM to 0.3 mM did not increase the GFP fluorescence in cultures at 37 °C. Therefore, for the cultures at 20 °C, only 0.1 and 0.2 mM IPTG were used. The OTR values in cultures at 20 °C were very low, compared to cultures at 37 °C (Fig. 2a and 3b). This may be due to the low oxygen consumption rate of the cells, as the growth at 20 °C was slow (some cultures lasted up to 50 h, Fig. 3a). The strains exhibited prolonged *lag* phases, lasting between 10 and 20 h (Fig. 3a). The GFP fluorescence increased parallel to cell growth. In cultures of the *rpoS* mutant, both the GFP fluorescence and biomass signals strongly increased immediately after the OTR decrease, indicating glucose exhaustion (Fig. 3b). This could be related to the downregulation of stationary phase genes in this mutant, which in turn may release cellular resources for further GFP expression. The specific growth rate was severely reduced in the *rpoD, rpoS* and *rpoH* mutants, compared to the WT (Fig. 3b). With exception of the *rpoN* mutant, all the other strains attained higher biomass than the WT (Fig. 3b). The maximal GFP fluorescence was similar in cultures of the WT and *rpoN* and *fliA* mutant strains, while for the *fecI* mutant, it was 20 % lower than for the WT under the two induction levels (Fig. 3b). In contrast GFP expression was remarkably higher in the *rpoD, rpoH* and *rpoS* mutant strains than in the WT (Fig. 3b). Particularly, the GFP fluorescence in cultures of the *rpoS* mutant was up to 19-fold higher than in cultures of the WT strain. Changing the IPTG concentration from 0.1 to 0.2 mM increased GFP fluorescence by 23 % in cultures of the *rpoS* mutant strain, whereas no relevant changes were observed for the other strains. The *rpoS* mutant exhibited the highest specific GFP fluorescence yield and fluorescence emission rate, which were up to 9 and 4-fold greater, respectively, than those of the WT strain.

**Figure 3.**
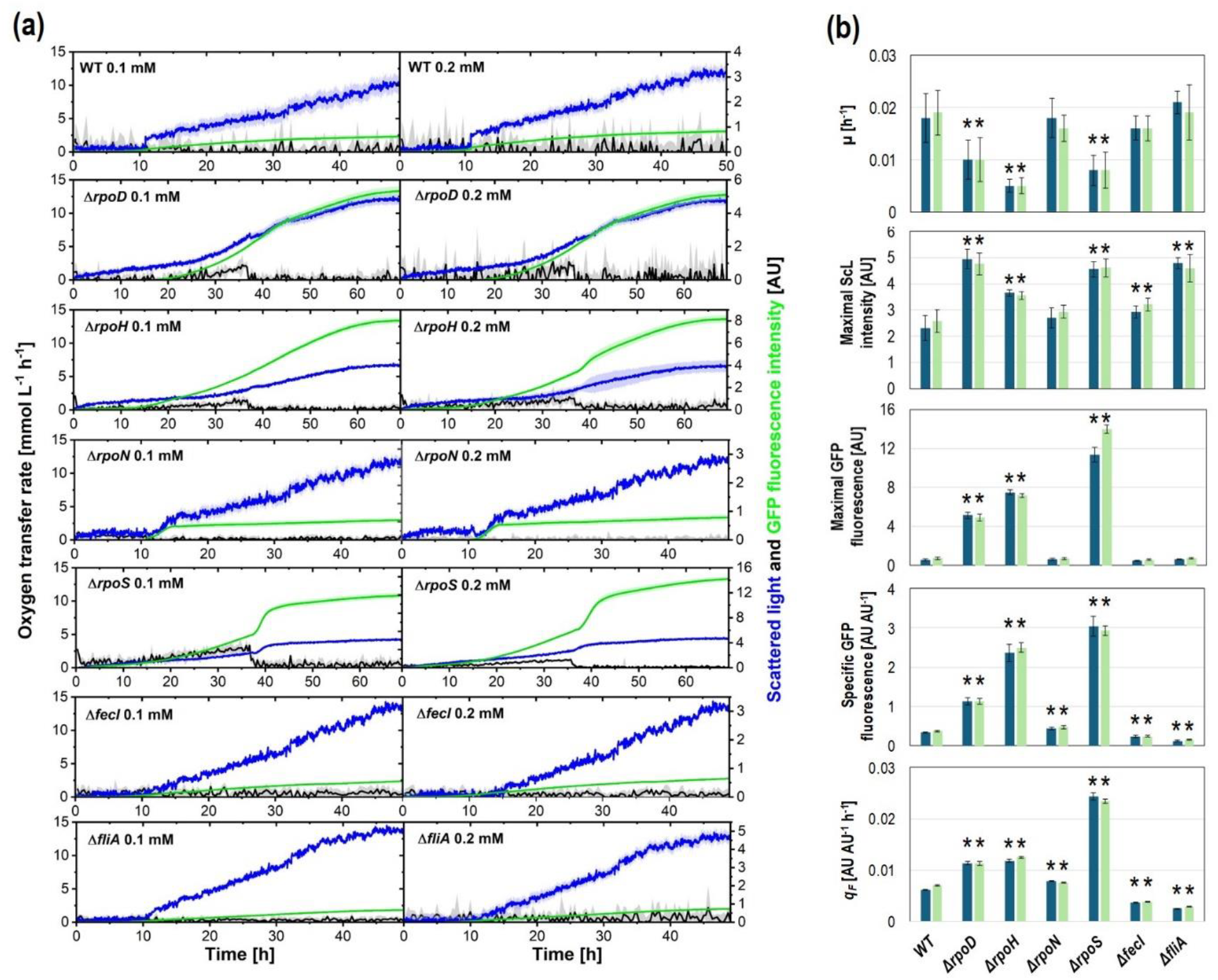
(a) Culture profile of the strains bearing the plasmid pV21 in mineral medium at 20 °C. GFP expression was induced with 0.1 or 0.2 mM IPTG from inoculation. Cell growth (blue lines) was monitored as scattered light (ScL) intensity. The oxygen transfer rate and GFP fluorescence were also monitored. Shaded bands indicate the standard deviation between replicates. (b) Main culture parameters using 0.1 (blue bars), or 0.2 (green bars) mM IPTG. Specific growth rate (*µ*), GFP fluorescence, and GFP fluorescence emission rate (*q*_*F*_) were calculated over the exponential growth phase. * Indicates significant difference (*n* =4; *p*<0.05), compared to the WT strain under the same conditions.

At 20 °C, *E. coli* cells experience a cold shock, which lowers cell growth and biomass yields, as can be seen comparing the results of Fig. 2 with those of Fig. 3. The sigma factor σ^S^ is essential for the response to cold shock, as it is involved in this stress response and activates genes that code for membrane proteins to strengthen them (White-Ziegler et al., 2008). A deletion of the *rpoS* gene leads to a lower growth of the cells (Fig. 3b), which are more sensitive to the cold shock due to the lack of membrane support. However, at 20 °C, the highest *q*_*F*_ can be observed for this mutant. The suppressed cold stress response appears to provide additional metabolic space for GFP expression. Therefore, the *rpoS* deletion seems to be an attractive option for improving recombinant protein expression at low temperatures in *E. coli*.

GFP expression was also evaluated in the widely used LB medium, at 37 °C and with 0.1 and 0.2 mM IPTG. The results are shown in Fig 4. No growth of the *rpoH* mutant was observed and consequently this strain is excluded from Fig. 4. The OTR time profile suggests the exhaustion of several nutrients during cell growth, which agrees with the complex composition of this culture medium (Fig. 4a). Like in previous experiments, the GFP fluorescence increased proportionally to the cell growth (Fig. 4a). The growth rate was lower in all mutants and induction levels, except for the *rpoN* mutant with 0.1 and 0.2 mM IPTG, and the *fecI* mutant with 0.1 mM IPTG (Fig. 4b). The individual deletion of the sigma factors resulted in lower attained biomass, except in the case of the *rpoN* mutant. The unaffected growth rate and biomass synthesis of the *rpoN* mutant suggest that the genes controlled by this sigma factor are less relevant in the rich LB than in the mineral media, in which the growth rate of the *rpoN* mutant was noticeably lower than that of the WT (Fig. 1-3). Furthermore, the biomass attained in cultures of the *rpoN* mutant was very similar to those of the WT, which contrasts with the rest of the mutant strains, that exhibited lower biomass values (Fig. 4b). In contrast, only the *rpoD, rpoS* and *fliA* mutants reached maximal GFP fluorescence values greater than the WT. During the exponential growth phase, the specific GFP fluorescence of the *fecI* and *fliA* mutants was approx. 20 % lower than the corresponding value for the WT. The *rpoN* mutant achieved a specific GFP fluorescence value 50 % higher than that of the WT. Remarkably, the specific GFP fluorescence of mutant strains *rpoS* and *rpoD* were approx. 3 and 9-fold greater than the achieved by the WT (Fig. 4b). Consequently, the specific GFP fluorescence emission rate of the mutant strain *rpoD* was approx. 7-fold higher, compared to the WT (Fig. 4b). In general, the accumulated oxygen transferred in cultures of the mutant strains was slightly but significantly (p<0.05) lower than that of the WT (Fig. 4b). The lowest value was observed in cultures of the mutant strain *rpoD*, which was approx. 20 % lower than that of cultures of the WT. The higher GFP expression and lower oxygen consumption render the *rpoD* mutant an excellent option for protein expression in a rich medium. A reason for the superior performance of this strain could be related to a lower proteomic burden in the rich LB medium, due to lesser expression of the proteins controlled by the transcriptional factor RpoD. Further studies are required to fine-tune the expression level of *rpoD* as a strategy to optimize recombinant protein expression in rich media.

**Figure 4.**
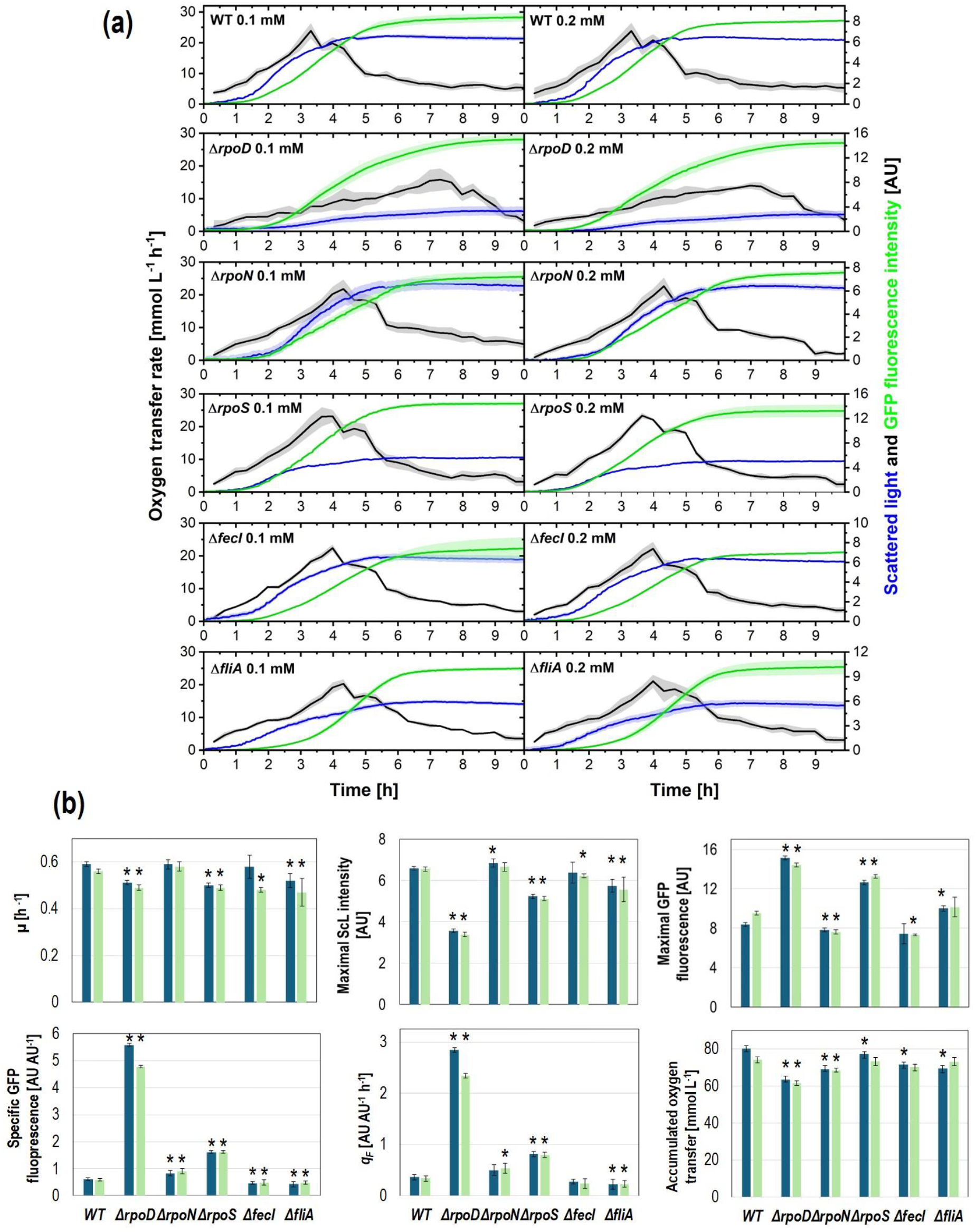
**Figure 3**. (a) Culture profile of the strains bearing the plasmid pV21 in LB medium at 37 °C. GFP expression was induced with 0.1 or 0.2 mM IPTG from inoculation. Cell growth was monitored as scattered light (ScL) intensity. The oxygen transfer rate and GFP fluorescence were also monitored. Shaded bands indicate the standard deviation between replicates. (b) Main culture parameters using 0.1 (blue bars), or 0.2 (green bars) mM IPTG. Specific growth rate (*µ*), GFP fluorescence, and GFP fluorescence emission rate (*q*_*F*_) were calculated over the exponential growth phase. * Indicates significant difference (*n* = 4; *p*<0.05), compared to the WT strain under the same conditions.

Taken together, the results show that deletion of sigma factors is a useful tool to improve the production of recombinant protein in *E. coli*, since they clearly affect the expression levels, growth rates and oxygen consumption. As this study shows, the best target for deletion will depend on the culture conditions like temperature and media, and should be decided depending on the application.

## 1 MATERIALS AND METHODS

### 1.1 Strains and plasmid

The K-12 derivative BW25113 *Escherichia coli* was used as wild type (WT). The mutans strains with individual interruptions of the genes *rpoD, rpoN, rpoS, fecI* and *fliA*, were obtained from the Keio Collection (Baba et al., 2006). The gene *rpoH* was interrupted using the method described by Datsenko and Wanner (2000), except that the mutant strain was incubated at 20 °C. The strains were transformed with the high copy number plasmid pV21, which contains the gene of the green fluorescence protein (GFP) under transcriptional control of the *lac* promoter, and a gene conferring spectinomycin resistance (Velazquez et al., 2022). Transformed and untransformed strains were plated in LB-agar in Petri dishes and grown at 37 °C for 14-18 h, except for the *rpoH* mutant, which was grown at 20 °C for 48 h. One colony per strain was selected and grown in Terrific Broth (TB) medium at 20 °C (for the *rpoH* mutant) or 37 °C (for all the other strains) until reaching an optical density at 600 nm (OD_600_) of approx. 6-8. Then, 0.9 mL of such culture broth was mixed with sterile solution of glycerol (80% v/v) and immediately frozen at -80 °C. The genotypes of the mutant strains were confirmed by PCR amplification of the involved gene using the oligomers described in the Supplementary File. To further confirm the PCR product of the Δ *rpoD* strain, 6 colonies were sampled and subjected to PCR amplification.

### 1.2 Media composition

All chemicals applied for media preparation were of analytical grade and purchased from Carl Roth GmbH, if not stated otherwise. The composition of LB was (in g L^-1^): yeast extract, 5; tryptone, 10; NaCl, 5. The composition of TB was (in g L_-1_): yeast extract, 24; tryptone, 20; glycerol, 4. The mineral medium composition was (in g L^-1^): (NH_4_)_2_SO_4_, 6.5; NH_4_Cl, 0.5; K_2_HPO_4_, 3.0; Na_2_SO_4_, 2.0; MgSO_4_ • 7H_2_O, 0.5; thiamine hydrochloride 0.01; MOPS, 41.85; plus 1 mL of trace elements solution per L of medium. Spectinomycin was purchased from Sigma Aldrich. The initial pH was adjusted to 7.4 by adding 0.2 M NaOH. The trace elements solution was prepared and autoclaved separately. The composition of the trace element solution was (in g L^-1^): ZnSO_4_• 7H_2_O, 0.54; CuSO_4_ • 5H_2_O, 0.48; MnSO_4_ • H_2_O, 0.30; CoCl_2_ • 6H_2_O, 0.54; FeCl_3_ • 6H_2_O, 41.76; CaCl_2_ • 2H_2_O, 1.98; Na_2_EDTA • 2H_2_O, 33.4. Glucose was added from a separately autoclaved solution at a concentration of 500 g L_-1_, to final concentrations of 4 g L_-1_ for cultures of the untransformed strains, or 3.5 g L_-1_ for cultures of the plasmid-bearing strains. For all cultures of plasmid bearing cells, the media were supplemented with 0.1 g L_-1_ spectinomycin. GFP expression was induced with isopropyl β-D-1-thiogalactopyranoside (IPTG) added prior to inoculation at final concentrations of 0.1, 0.2 or 0.3 mM.

### 1.3 Microbioreactor cultures

The strains were characterized in microbioreactor cultures in lysogeny broth (LB) or in the mineral medium supplemented with 4 g L^-1^ glucose. For cultures in LB, a preculture was performed by transferring 0.05 mL of the cryopreserved cells to 8 mL of LB medium contained in 250 mL Erlenmeyer flask. For cultures in the mineral medium, 0.05 mL of the cryopreserved cells were transferred to 8 mL of TB medium contained in 250 mL Erlenmeyer flask. The cells were grown at 20 (for the *rpoH* mutant) or 37 °C (for all other strains), and 350 rpm in an orbital shaker of 50 mm shaking diameter for 6 h. For the main cultures in mineral medium, 0.05 mL from the first preculture in TB medium were transferred to 8 mL of mineral medium contained in a 250 mL Erlenmeyer flask and cultured at 20 (for the *rpoH* mutant) or 37 °C (for all other strains), and 350 rpm in an orbital shaker of 50 mm shaking diameter until mid-exponential growth phase (approx. 14-16 h). The obtained cells were washed with 0.9 % sterile NaCl solution and used to inoculate the main cultures. Microbioreactor cultures were carried out in 48-round transparent bottom wells microtiter plates (MTP) (Beckman Coulter, Krefeld, Germany) with a liquid volume of 0.75 mL per well, an initial OD_600_ of 0.1 units, at 20 or 37 °C, 1000 rpm and 3 mm shaking diameter (Climo-Shaker ISF1-X, Kuhner, Birsfelden, Switzerland). The shaken monitoring system, named µRAMOS-BioLector combination, is in-house constructed, and enables measurement of the OTR in every individual well of the MTP based on the oxygen partial pressure and oxygen-dependent emission of fluorescence sensors (Flitsch et al., 2016), and synchronized online measurement of fluorescence and scattered light (Ladner et al., 2016). The MTPs were covered with a gas-permeable polyolefin sealing foil (HJ-Bioanalytik GmbH, Erkelenz, Germany), to reduce evaporation and prevent contaminations. Cell growth was monitored by measuring the scattered light (ScL) intensity at a wavelength of 620 nm. The wavelengths for excitation and fluorescence emission of GFP were 480 and 507 nm, while for NADH were 340 and 460 nm, respectively.

### 1.4 *rpoD* gene transcription level

Triplicate cultures of the WT and *rpoD* mutant strains were carried out in the mineral medium at 37 °C as explained above. Samples were taken during the exponential growth of the cells. RNA extraction, purification, RT-qPCR conditions, and relative transcription level calculation are described in detail elsewhere (Aguilar et al., 2012).

### 1.5 Data analysis

The data from fluorescence and scattered light (ScL) intensities are represented as the corresponding reading minus the lowest reading (usually obtained during the first 20 min of culture). The specific fluorescence (SF) was calculated as the mean of the fluorescence divided by the ScL readings in every time point during the period involved. Specific yields were calculated by the proper mass balances over the time periods involved. The specific fluorescence emission rate was calculated by multiplying the specific growth rate by the specific GFP fluorescence intensity. Accumulated oxygen transfer was calculated by integration of the area under the curve in the plot of OTR versus time using the Integration gadget of the Origin software (OriginLab Corporation). For comparison between two groups means, unpaired Student’s *t*-tests were performed.

## AUTHOR CONTRIBUTIONS

*Conceptualization*: A. R. Lara, G. Gosset, J. Büchs. *Experimental design, planning, execution, and data acquisition and analysis*: L. Pohlen, E. García, N. Flores, L. M. Martínez, A. R. Lara, G. Gosset. *Funding acquisition*: A. R. Lara, G. Gosset, J. Büchs. *Writing and review of manuscript*: all authors.

## ACKNOWLEDGMENTS

This work was supported by CONACyT grant A1-S-8646.

## DATA AVAILABILITY STATEMENT

The data that support the findings of this study are available from the corresponding authors upon reasonable request.

## References

Aguilar, C., Escalante, A., Flores, N., de Anda, R., Riveros-McKay, F., Gosset, G., Morett, E., & Bolívar, F. (2012). Genetic changes during a laboratory adaptive evolution process that allowed fast growth in glucose to an Escherichia coli strain lacking the major glucose transport system. BMC Genomics, 13, 385. 10.1186/1471-2164-13-385

Baba, T., Ara, T., Hasegawa, M., Takai, Y., Okumura, Y., Baba, M., Datsenko, K. A., Tomita, M., Wanner, B. L., & Mori, H. (2006). Construction of Escherichia coli K-12 in-frame, single-gene knockout mutants: the Keio collection. Molecular Systems Biology, 2, 2006.0008.

Bafna-Rührer, J., Bhutada, Y. D., Orth, J. V., Øzmerih, S., Yang, L., Zielinski, D., & Sudarsan, S. (2024). Repeated glucose oscillations in high cell-density cultures influence stress-related functions of Escherichia coli. PNAS Nexus, 3(9), pgae376. 10.1093/pnasnexus/pgae376

Cho, B. K., Kim, D., Knight, E. M., Zengler, K., & Palsson, B. O. (2014). Genome-scale reconstruction of the sigma factor network in Escherichia coli: topology and functional states. BMC Biology, 12, 4. 10.1186/1741-7007-12-4

Chou, C. H., Bennett, G. N., & San, K. Y. (1996). Genetic manipulation of stationary-phase genes to enhance recombinant protein production in Escherichia coli. Biotechnology and Bioengineering, 50(6), 636–642. 10.1002/(SICI)1097-0290(19960620)50:6<636::AID-BIT4>3.0.CO;2-L

Datsenko, K. A., & Wanner, B. L. (2000). One-step inactivation of chromosomal genes in Escherichia coli K-12 using PCR products. Proceedings of the National Academy of Sciences of the United States of America, 97(12), 6640–6645. 10.1073/pnas.120163297

Dong, T., & Schellhorn, H. E. (2009). Control of RpoS in global gene expression of Escherichia coli in minimal media. Molecular Genetics and Genomics: MGG, 281(1), 19–33. 10.1007/s00438-008-0389-3

Flitsch, D., Krabbe, S., Ladner, T., Beckers, T., Schilling, J., Mahr, S., Conrath, U., Schomburg W.K. & Büchs, J. (2016). Respiration activity monitoring system for any individual well of a 48-well microtiter plate. Journal of Biological Engineering, 10, 14. 10.1186/s13036-016-0034-3

Goodall, E. C. A., Robinson, A., Johnston, I. G., Jabbari, S., Turner, K. A., Cunningham, A. F., Lund, P. A., Cole, J. A., & Henderson, I. R. (2018). The Essential Genome of Escherichia coli K-12. mBio, 9(1), e02096–17. 10.1128/mBio.02096-17

Han, J. H., Jung, S. T., & Oh, M. K. (2021). Improved Yield of Recombinant Protein via Flagella Regulator Deletion in Escherichia coli. Frontiers in Microbiology, 12, 655072. 10.3389/fmicb.2021.655072

Ladner, T., Held, M., Flitsch, D., Beckers, M., & Büchs, J. (2016). Quasi-continuous parallel online scattered light, fluorescence and dissolved oxygen tension measurement combined with monitoring of the oxygen transfer rate in each well of a shaken microtiter plate. Microbial Cell Factories, 15(1), 206. 10.1186/s12934-016-0608-2

Ladner T., Mühlmann M., Schulte A., Wandrey G., & Büchs J. (2017). Prediction of Escherichia coli expression performance in microtiter plates by analyzing only the temporal development of scattered light during culture. Journal of Biological Engineering, 11:20.

Lara, A. R., Utrilla, J., Martínez, L. M., Krausch, N., Kaspersetz, L., Hidalgo, D., Cruz-Bournazou, N., Neubauer, P., Sigala, J. C., Gosset, G., & Büchs, J. (2024). Recombinant protein expression in proteome-reduced cells under aerobic and oxygen-limited regimes. Biotechnology and Bioengineering, 121(4):1216–1230. 10.1002/bit.28645

Lin, T. H., Cheng, S. Y., Lin, Y. F., & Chen, P. T. (2024) Development of the Low-Temperature Inducible System for Recombinant Protein Production in Escherichia coli Nissle 191. Journal of Agricultural and Food Chemistry, 72(13), 7318–7325. 10.1021/acs.jafc.4c01075

Mühlmann, M. J., Forsten, E., Noack, S., & Büchs, J. (2018). Prediction of recombinant protein production by Escherichia coli derived online from indicators of metabolic burden. Biotechnology Progress, 34(6), 1543– 1552. 10.1002/btpr.2704

Reitzer, L., & Schneider, B. L. (2001). Metabolic context and possible physiological themes of sigma(54)-dependent genes in Escherichia coli. Microbiology and Molecular Biology Reviews: MMBR, 65(3), 422–444. 10.1128/MMBR.65.3.422-444.2001

San-Miguel, T., Pérez-Bermúdez, P., & Gavidia, I. (2013). Production of soluble eukaryotic recombinant proteins in E. coli is favoured in early log-phase cultures induced at low temperature. Springerplus. 2(1):89. 10.1186/2193-1801-2-89.

Schavemaker, P. E., & Lynch, M. (2022). Flagellar energy costs across the tree of life. eLife, 11, e77266. 10.7554/eLife.77266

Shimada, T., Yamazaki, Y., Tanaka, K., & Ishihama, A. (2014). The whole set of constitutive promoters recognized by RNA polymerase RpoD holoenzyme of Escherichia coli. PloS One, 9(3), e90447. 10.1371/journal.pone.0090447

Velazquez, D., Sigala, J. C., Martínez, L. M., Gaytán, P., Gosset, G., & Lara, A. R. (2022). Glucose transport engineering allows mimicking fed-batch performance in batch mode and selection of superior producer strains. Microbial Cell Factories, 21(1), 183. 10.1186/s12934-022-01906-1

White-Ziegler, C. A., Um, S., Pérez, N. M., Berns, A. L., Malhowski, A. J., & Young, S. (2008). Low temperature (23 degrees C) increases expression of biofilm-, cold-shock- and RpoS-dependent genes in Escherichia coli K-12. Microbiology (Reading, England), 154(1), 148–166. 10.1099/mic.0.2007/012021-0

Yamamoto, N., Nakahigashi, K., Nakamichi, T., Yoshino, M., Takai, Y., Touda, Y., Furubayashi, A., Kinjyo, S., Dose, H., Hasegawa, M., Datsenko, K. A., Nakayashiki, T., Tomita, M., Wanner, B. L., & Mori, H. (2009). Update on the Keio collection of Escherichia coli single-gene deletion mutants. Molecular Systems Biology, 5, 335.

Ziegler, M., Zieringer, J., Döring, C. L., Paul, L., Schaal, C., & Takors, R. (2021). Engineering of a robust Escherichia coli chassis and exploitation for large-scale production processes. Metabolic Engineering, 67, 75–87. 10.1016/j.ymben.2021.05.011

