## Supplementary material for "Sigma factors as potential targets to enhance recombinant protein expression"

**Table S1.** Primer and expected PCR product size for each gene in the WT and corresponding mutant.

| Gene | Primers | Expected PCR product length in WT strain | Expected PCR product length in mutant strain |
|---|---|---|---|
| *rpoD* | Forward 5´- TGG TTT AAG CAA CGA AGA ACG -3´<br>Reverse 5´- CGT TTT TTA TCG CCC ACG CAC -3´ | 1980 | 1458 |
| *fecI* | Forward 5´- CAC GGC ATA CCA GTG GGA AG -3´<br>Reverse 5´- CAG AAG CAG AGG CTC CAT CC -3´ | 930 | 1731 |
| *rpoN* | Forward 5´- GCG CTT ATA TCG TCA GTC A -3´<br>Reverse 5´- AGT GTT GCA TCT GAG GTG -3´ | 1775 | 1661 |
| *fliA* | Forward 5´- TCG CGA ACT AAG GTA ATG C -3´<br>Reverse 5´- ACC TGA TTA ACT GAG ACT G -3´ | 1197 | 1773 |
| *rpoS* | Forward 5´- GGC TTG GTG CGT CAC ATA T -3´<br>Reverse 5´- CCA GTT CAA CAC GCT TGC A -3 | 1387 | 1751 |

**(a)**

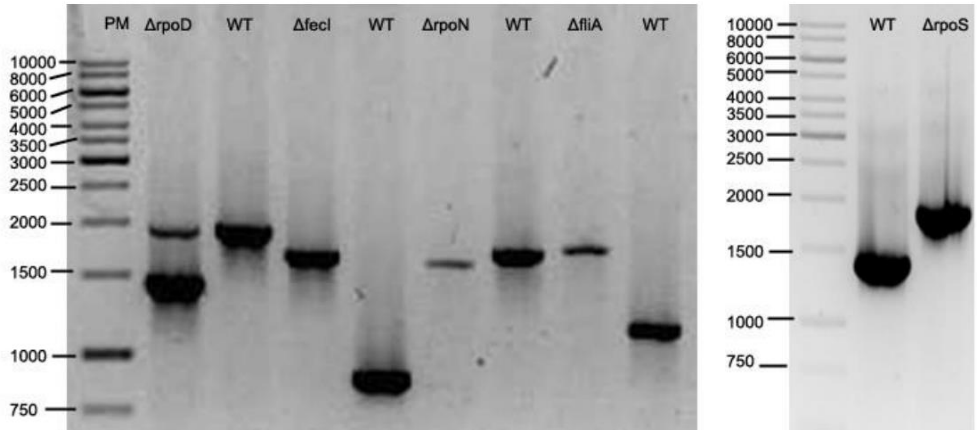


**(b)**

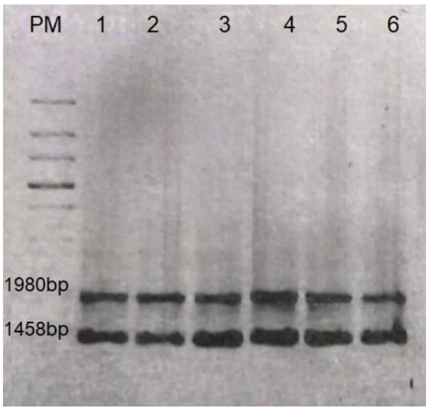


**Figure S1.** PCR amplification of the target genes in the different mutants. (a): Agarose gel image showing the obtained bands and corresponding weights. (b): PCR amplification of *rpoD* in 6 different colonies. PM: molecular weight ladder.

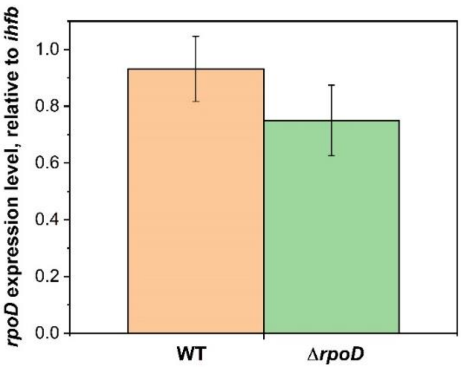


**Figure S2.** *rpoD* expression level relative to *ihfB* in the WT and *rpoD* mutant strains. Vertical bars indicate the standard deviation between replicates ($n$ = 7)
